# Brain morphometric changes in congenitally blind subjects: a 7 Tesla MRI study

**DOI:** 10.1101/2022.05.18.492367

**Authors:** Ron Kupers, Minye Zhan, Samuel Paré, Laurence Dricot, Maarten Vaessen, Beatrice de Gelder

## Abstract

We used ultra-high field (7 Tesla) magnetic resonance imaging (MRI) at submillimeter resolution to assess structural brain changes in congenitally blind (CB) compared to matched normal sighted control (SC) subject groups. Region-of-interest analysis revealed grey matter (GM) volumetric reductions in the CB group in left cuneus and occipital pole, right posterior collateral sulcus and right occipito-temporal medial lingual sulcus. Non-visual areas with GM reductions in CB included the left central, postcentral and superior frontal gyri, and the right subcallosal gyrus. In contrast, there were no significant group differences in cortical thickness when using stringent statistical criteria. Regional differences in white matter (WM) showed an overall pattern similar as that of GM changes, characterized by volume reductions in occipital, parietal and temporal areas, but with additional reductions in precuneus and medial orbitofrontal cortex. Differences in cortical curvature were mostly situated in the occipital cortex and bore a close relationship with areas showing GM alterations; they may be indicative of increased cortico-cortical connectivity of the visually-deprived occipital cortex. The CB group had GM reductions in the basal ganglia, i.e., caudate nucleus, putamen, nucleus accumbens, globus pallidus, and thalamus. Within the cerebellum, GM and WM volumes were also reduced in the CB. Segmentation of the thalamus, hippocampus and amygdala into anatomic divisions revealed GM reductions in a number of thalamic nuclei, a few hippocampal regions, but not within amygdala. There were no findings of increased volume or cortical thickness in the CB group. Together, these data reveal a multitude of GM and WM reductions in CB, comprising not only the occipital cortex, but also temporal, parietal, and prefrontal cortices, as well as the basal ganglia and cerebellum. These findings in the CB may seem at odds with the large literature showing that the visually-deprived occipital cortex becomes a multimodal cortex responding to diverse non-visual sensory and cognitive inputs. The seeming mismatch between morphological atrophy and enhanced multimodality of occipital areas combined with superior performance by the CB in various non-visual tasks poses a challenge for our understanding of brain plasticity.

## Introduction

For humans and non-human primates, the visual system is by far the most important sense for perceiving the outside world. Anatomically, this primacy of the visual sense is reflected by the large cortical territory occupied by the primate visual system, whereby one third of the cortical mantle is involved in the processing of visual information.^1^ Congenital blindness (CB), or the loss of vision during post-natal life, must therefore have a dramatic influence on the structural and functional (re)organization of visual cortical areas. Indeed, there is ample evidence that the deprived human visual cortex remains metabolically active, responds to various forms of non-visual sensory input, and even assumes a role in higher cognitive functions such as memory and language processing.^2-5^ At the behavioral level, these changes are associated with an increased sensitivity of the remaining intact senses.^4^ Taken together, these observations suggest that blindness may promote a functional and anatomic restructuring of the primate brain.

Using volumetric magnetic resonance imaging (MRI) we previously showed that CB is associated with various alterations in grey matter (GM) and white matter (WM), both within and outside the occipital cortex.^6-8^ Although the most important anatomic findings in CB were observed in striate and extrastriate cortical areas, other brain areas also undergo alterations, including the basal ganglia^6^, somatosensory cortex,^9-14^ hippocampus,^7,15^ olfactory bulb^16^ and prefrontal cortex.^6^ Although there is a general agreement that CB is associated with relative volumetric reductions in striate and extrastriate cortices, there is still debate about the extent and type of structural changes outside the visual cortex. For instance, whereas some authors have reported volumetric increases in the post-central sulcus^10^, others found a decrease in this area^9,13^ or no change at all. The reasons for these discrepancies are unclear, but may reflect the low number of participants included in some studies, and the admixture of data from CB and early blind subjects. In addition, few studies have looked for differences within divisions of subcortical structures and cerebellum. For instance, while the majority of the studies have reported no volumetric change for the hippocampus as a whole, its segmentation revealed volumetric reduction in the posterior region^7^, or increases in the anterior division.^15^ There is hence a need for a more refined assessment of structural changes within subcortical GM structures occurring in association with CB.

The aim of the present study was to use ultra-high field MRI imaging at submillimeter spatial resolution in a homogenous group of CB subjects in order to obtain an accurate and exhaustive description of anatomical differences relative to a matched control group. We measured and quantified overall and regional group differences in GM and WM volumes, in cortical thickness, and cortical curvature, while using state-of-the-art tools to segment the thalamus, hippocampus, amygdala and cerebellum into their constituent nuclei or lobules.

## Materials and methods

### Participants

Twelve congenitally blind (CB; 5 females; age: 46.0±10.3 y, range: 29-62 y) and 11 matched sighted controls (SC; 4 females; age: 44.8±11.6 y, range: 25-61 y) participated in the study. All CB participants were blind from birth and had no residual visual perception. Demographic data are described in Supplementary Table 1.

### MRI data acquisition

MRI data were acquired at the Scannexus facility of Maastricht University (The Netherlands), using a 7T Magnetom scanner (Siemens, Erlangen, Germany), with a 1Tx/32Rx head coil (Nova Medical Inc., USA). For each participant, we collected T1-weighted, proton density (PD), and T2*-weighted image contrasts. The imaging parameters of the T1-weighted MPRAGE sequence were as follows: TR=3100 ms, TE=2.52 ms, TI=1500 ms, flip angle=5°, FOV read=230 mm, matrix size =384×384×256, resolution=0.6×0.599×0.599 mm, pixel bandwidth=180 Hz/Px, GRAPPA acceleration factor=3. The proton density sequence had the following parameters: MPRAGE, TR=1440 ms, TE=2.52 ms, flip angle=5°, FOV read=230 mm, matrix size=384×384×256, resolution=0.6×0.599×0.599 mm, pixel bandwidth=180 Hz/Px, GRAPPA acceleration factor=3. For the T2*-weighted contrast, we used a TurboFLASH sequence with TR=4910 ms, TE=16 ms, flip angle=5°, FOV read=230 mm, matrix size=384×384×256, 0.6×0.599×0.599 mm resolution, pixel bandwidth=480 Hz/Px, GRAPPA acceleration factor=3. The whole scanning session lasted approximately 60 minutes.

### MRI data processing

Data were preprocessed in BrainVoyager (BV version 22.0, Brain Innovation, The Netherlands). The T1 and PD images were first resampled to 0.6×0.6×0.6 mm isovoxels. To correct for inhomogeneities in image intensity, the T1 image was divided by the PD image. Thereafter, we removed background noise, skull-stripped the T1 image, and converted it into niftii format. The whole brain was then segmented using the FreeSurfer (version 7.2) image analysis pipeline (http://surfer.nmr.mgh.harvard.edu/). We used recon-all with the “highres” flag for data with native submillimeter resolution. The final segmentation (i.e., morphometry and surface data) was based on a subject-independent probabilistic atlas and subject-specific measured values. After passage through the pipeline, each segmentation was visually inspected and corrected, which was necessary in a few instances where small skull strips errors were found. The final data were resampled to an average brain template (fsaverage) and smoothed (FWHM=10 mm). We tested for group differences in cortical and subcortical grey matter volumes, cortical thickness, cortical gyrification (curvature) and white matter volume, using an atlas-based approach. ROI measures (atlas-based) were exported for statistical analysis into SPSS software (see below). We used the white-matter (wm.mgz), cortical and sub-cortical (aseg.mgz) segmentations as implemented in FreeSurfer. For cortical surface and thickness measures, we used the Destrieux atlas,^17^ whereas for thalamus, amygdala and hippocampus we used FreeSurfer plugins for subfields segmentation.^18,19,20^ The cerebellum was segmented into ten subfields (lobules) and WM, using CERES.^21^ We adjusted all the between-group comparisons by brain region for age, sex and intracranial volume, and corrected for multiple comparisons using the Benjamini–Hochberg procedure for FDR correction (at *Q=*0.05) in each of the atlases separately.

### Data availability

The data that support the findings of this study are available from the corresponding author, upon reasonable request.

## Results

### Volumetric changes

#### Global volumetric changes

As shown in Fig. 1, CB subjects had lower overall cortical volume (CB: 462,260± 50,947 mm^3^ vs. SC: 525,287±48,349 mm^3^; *P*=0.002), total subcortical gray matter (CB: 52,600±7,062 mm^3^ vs. SC: 58,923±3,359 mm^3^; *P*=0.005), cerebellar cortex (CB: 102,477±6,681 mm^3^ vs. SC: 109,769±4,099 mm^3^; *P*=0.0024) and cerebellar white matter (CB: 24,404±2,951 mm^3^ vs. SC: 28,874±3,725 mm^3^; *P*=0.0025). Subcortical structures in which CB subjects had reduced volume included bilateral thalamus (CB: 6,643±892 mm^3^ vs. SC: 7,568±861 mm^3^; *P*=0.0012), caudate nucleus (CB: 3,471±512 mm^3^; SC: 3,938±344 mm^3^; *P*=0.0012), putamen (CB: 4,768±847 mm^3^ vs. SC: 5,128±423 mm^3^; *P*=0.037), globus pallidus (CB: 1,736±281 mm^3^ vs. SC: 2,043±171 mm^3^; *P*=0.0001), brainstem (CB: 17,867±2,466 mm^3^ vs. SC: 20,204±1,675 mm^3^; *P*=0.0193), nucleus accumbens (CB: 542±107 mm^3^ vs. SC: 647±65 mm^3^; *P*=0.0004) and ventral diencephalon (CB: 3,637±416 mm^3^ vs. SC: 4,176±304 mm^3^; *P*=1.8957E-05) (Fig. 2). No significant group differences were found for whole volume measures of amygdala and hippocampus. For WM, CB subjects had lower WM volume for total brain (CB: 387,921±7,1124 mm^3^ vs. SC: 455,929±62,900 mm^3^; *P*=0.0025), total cerebellar WM (CB: 12,203±1,600 mm^3^ vs. SC: 14,437±2,208 mm^3^, *P*=0.0049), posterior part of corpus callosum (CB: 520±74 mm^3^ vs. SC: 633±118 mm^3^; *P*=0.0072) and optic chiasm (CB: 104±36 mm^3^ vs. SC: 151±19 mm^3^; *P*=0.0008). For none of the GM or WM structures did CB subjects have larger volumes compared to the SC group.

**Figure 1:**
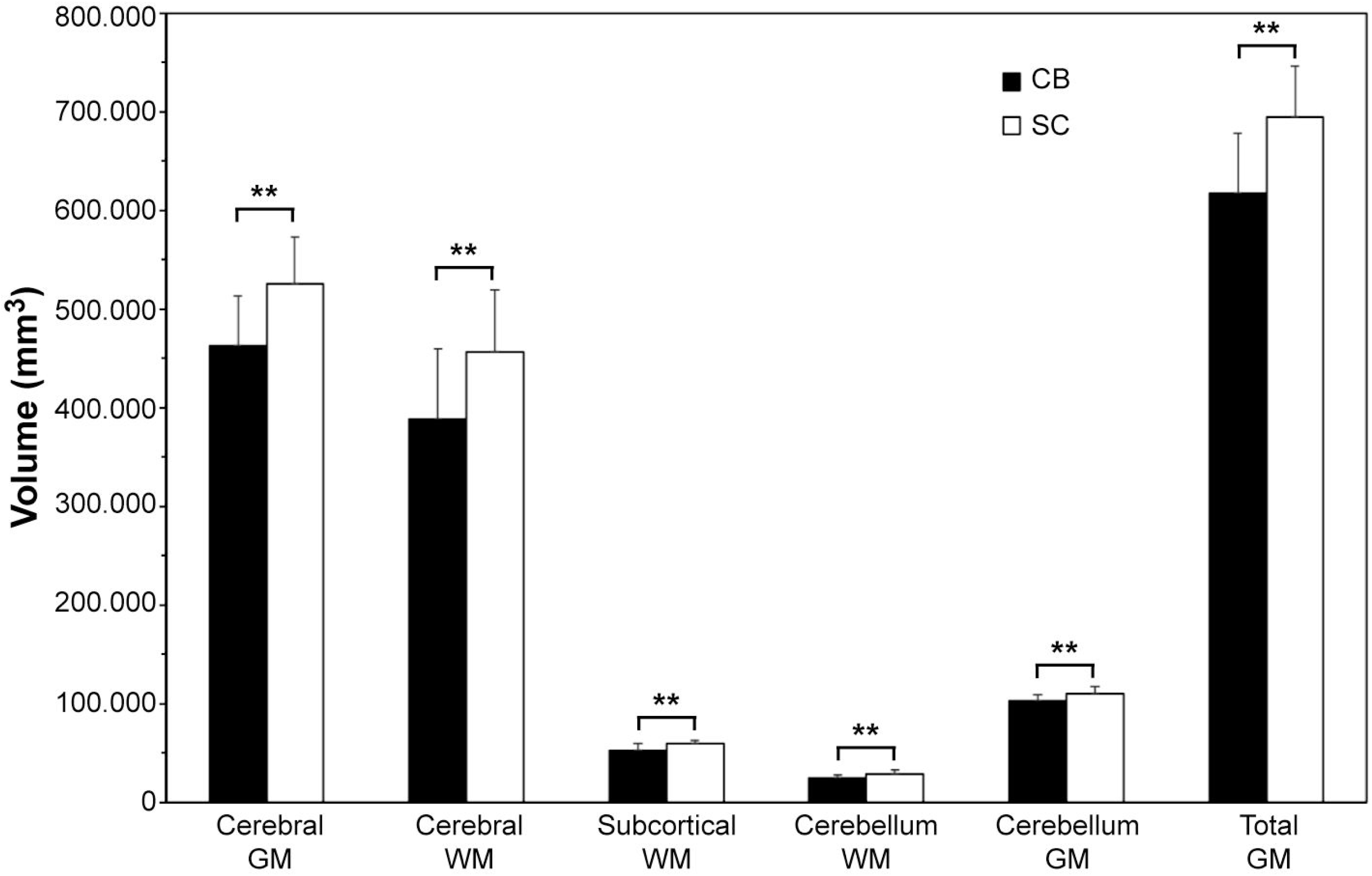
Alterations in global measures of cerebral, subcortical and cerebellar GM and WM in CB subjects. Abbreviations: CB: congenitally blind; SC: sighted controls; GM: grey matter; WM: white matter. **: P<0.01 (FDR-corrected)

**Figure 2:**
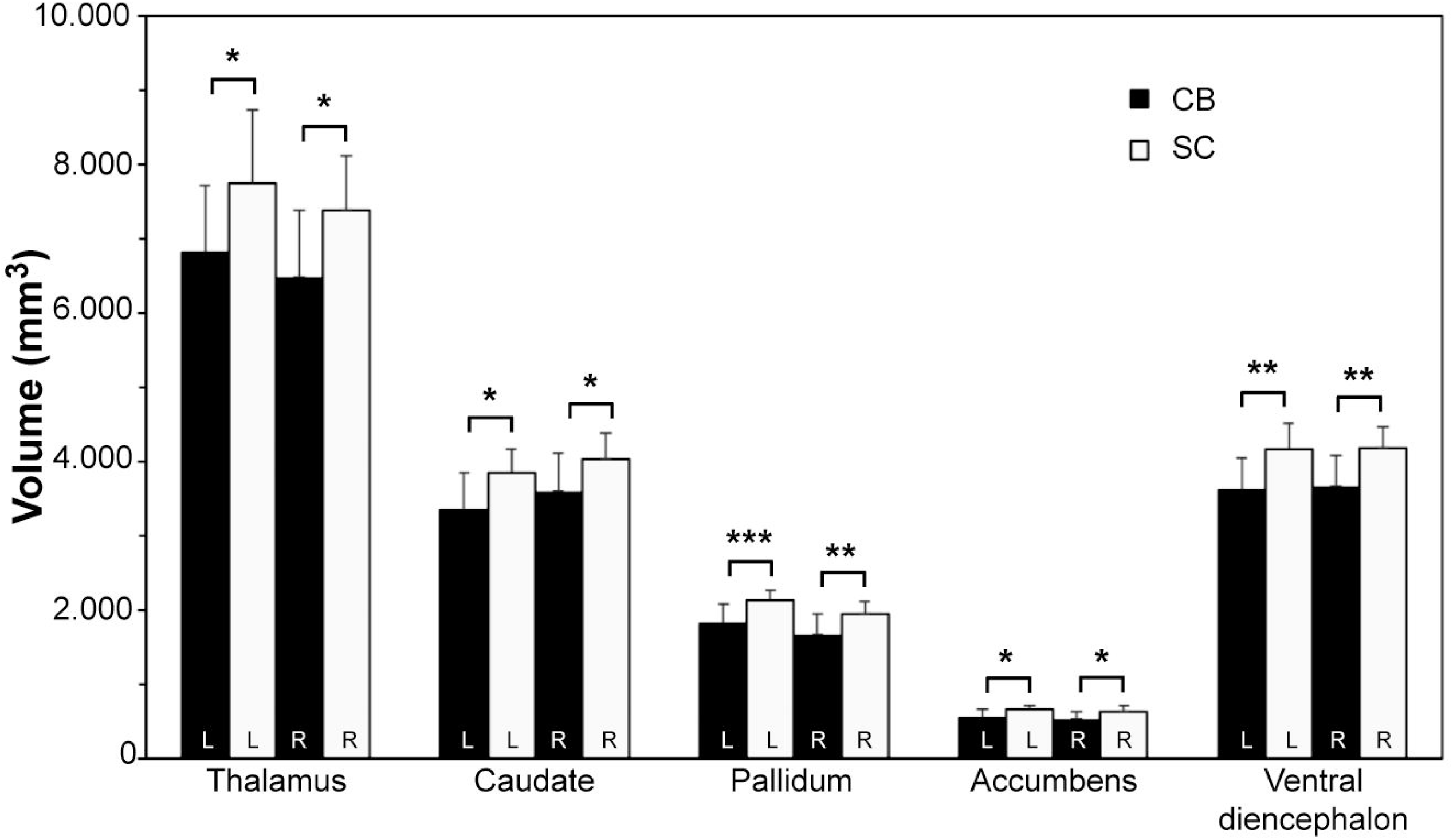
Volumetric reductions in subcortical structures in CB subjects. Abbreviations: CB: congenitally blind; SC: sighted controls; L: left; R: right. *: P<0.05 (FDR-corrected); **: P<0,01 (FDR-corrected), ***: P<0.001 (FDR-corrected).

### Atlas-based volumetric changes

#### Cortical grey matter volume

All significant group differences went in the direction of volumetric reductions in the CB group (Fig. 3). Within visual cortical areas, CB participants had lower volume in left cuneus (CB: 2,865±374 mm^3^ vs. SC: 4,180±838 mm^3^; *P*= 0,0001), left occipital pole (CB: 2,754±471 mm^3^ vs. SC: 3,470±435 mm^3^; *P*=0.0021), right posterior transverse collateral sulcus (CB: 482±99 mm^3^ vs. SC: 725±100 mm^3^; *P*=1.96E-05), and right occipito-temporal medial lingual sulcus (CB: 4,753±687 mm^3^ vs. SC: 5,617±1158 mm^3^; *P*=0.0006). The largest relative reductions were in the left cuneus (31%) and right posterior transverse collateral sulcus (34%). When using a more lenient statistical threshold (*P*≤0.05, non-FDR-corr.), we found additional volumetric reductions in other visual areas, including the right cuneus (CB: 3,250±391 mm^3^ vs. SC: 4,062±952 mm^3^; *P*=0.011), left calcarine sulcus (CB: 2,236±542 mm^3^ vs. SC: 3,154±821 mm^3^; *P*=0.005), left superior occipital sulcus and transverse occipital sulcus (CB: 1,419 ± 240 mm^3^ vs. SC: 1,803±397 mm^3^; *P*=0.011), and right parieto-occipital sulcus (CB: 2,615±518 mm^3^ vs. SC: 3,269±596 mm^3^; *P*=0.0072). CB subjects also showed significant volumetric reductions in parietal and prefrontal cortical areas, including left central sulcus (CB: 3,548±358 mm^3^ vs. SC: 4,069±393 mm^3^; *P*=0.0016), left postcentral gyrus (CB: 4,111±713 mm^3^ vs. SC: 5,192±875 mm^3^; *P*=0.0021), left superior frontal gyrus (CB: 4,172±803 mm^3^ vs. SC: 5,264±918 mm^3^; *P*=0.0012), and right subcallosal gyrus (CB: 888±220 mm^3^ vs. SC: 1,178±122 mm^3^; *P*=0.00025). At the more lenient statistical threshold (non-FDR-corr.), additional volumetric reductions were observed in the right postcentral gyrus (CB: 3,902±616 mm^3^ vs. SC: 4,711±656 mm^3^; *P*=0.0072), left precentral gyrus (CB: 5,867±655 mm^3^ vs. SC: 6,859±881 mm^3^; P=0.0035), inferior part of the left precentral sulcus (CB: 2,359±792 mm^3^ vs. SC: 3,044±695 mm^3^; *P*=0,0054), right angular gyrus (CB: 6,959±1,798 mm^3^ vs. SC: 8,876±1,308 mm^3^; *P*=0.0058) and right intraparietal sulcus and transverse parietal sulci (CB: 3,886±933 mm^3^ vs. SC: 4,955±769 mm^3^; *P*=0.0036).

**Figure 3:**
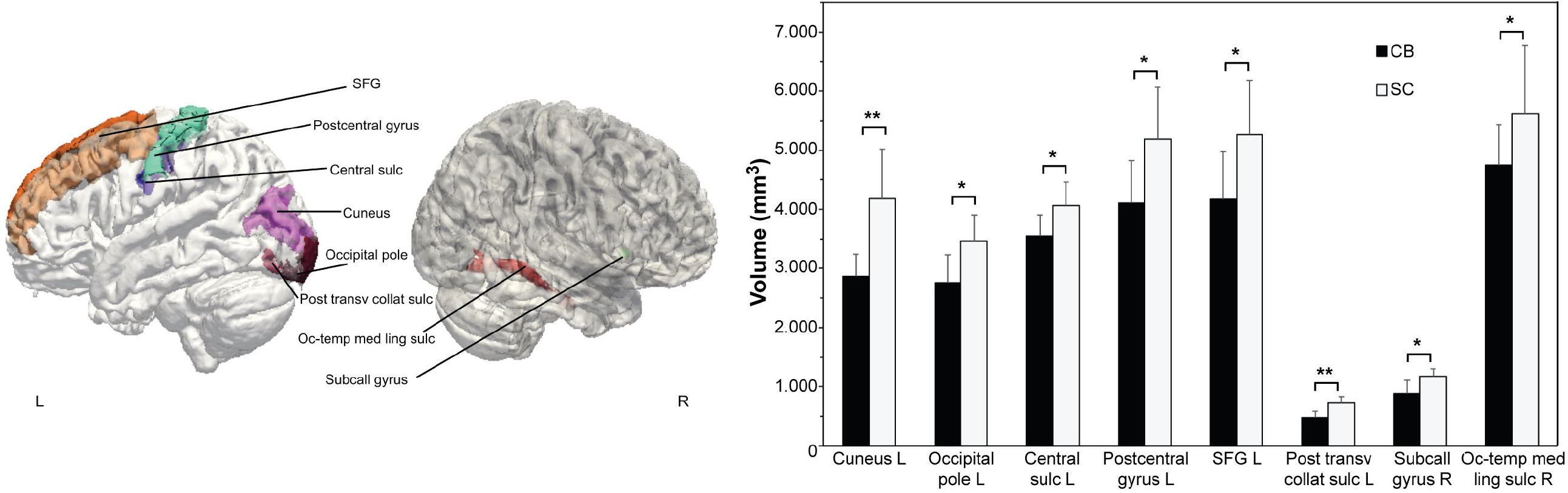
Alterations in cortical surface volume in CB subjects. ROI analysis using the Destrieux brain atlas. The left part of the figure shows the GM ROIs with significant changes projected on a brain template. Abbreviations: SFG: superior frontal gyrus; Post. transv. collat. sulc: posterior transverse collateral sulcus; Subcall: subcallosal; Oc-temp med ling sulc: lingual part of the medial occipito-temporal gyrus. *: P<0.05 (FDR-corrected); **: P<0.01 (FDR-corrected).

### White matter volume

The CB group showed widespread and substantial reductions in WM volumes comparted to the sighted controls in occipital, parietal, temporal and prefrontal areas (Fig. 4). In occipital areas, WM volume was reduced in bilateral cuneus (left: CB: 1,499±329 mm^3^ vs. SC: 2,244±539 mm^3^; *P*=0.0008; right: CB: 1,728±314 mm^3^ vs. SC: 2,263±492 mm^3^; *P*=0.0038), bilateral pericalcarine (left: CB: 1,823±536 mm^3^ vs. SC: 3,050±948 mm^3^; *P*=0.0030; right: CB: 2,026±526 mm^3^ vs. SC: 2,949±890 mm^3^; *P*=0.0083), bilateral lateral occipital (left: CB: 6,869±1,296 mm^3^ vs. SC: 9,219±1,451 mm^3^; *P*=0.0023; right: CB: 7,066±1,393 mm^3^ vs. SC: 9,021±1,620 mm^3^; *P*= 0.0098), bilateral lingual (left: CB: 3,481±972 mm^3^ vs. SC: 4,600±869 mm^3^; *P*=0.011; right: CB: 3,645±681 mm^3^ vs. SC: 4,908±1,161 mm^3^; P=0.0061), and right fusiform (CB: 4,938±1,091 mm^3^ vs. SC: 6,433±1,044 mm^3^; *P*=0.0015). The largest relative decreases in volume were in cuneus (left: -33%, right: -24%) and the pericalcarine area (left: -40%, right: -31%); volumes in other occipital areas were 22-26% lower in the CB group. In parietal areas, CB subjects had lower WM volumes in left precentral (CB: 11,206±1,658 mm^3^ vs. SC: 12,863±1,467 mm^3^; *P*=0.0068), bilateral postcentral (left: CB: 6,341±1,108 mm^3^ vs. SC: 7,796±1,763 mm^3^; *P*=0.0091; right: CB: 6,292±1155 mm^3^ vs. SC: 7,700±1802 mm^3^; *P*=0.014), left paracentral (CB: 3,132±457 mm^3^ vs. SC: 3,765±505 mm^3^; *P*=0.0076), right superior parietal (CB: 9,221±2,107 mm^3^ vs. SC: 10,985±1,644 mm^3^; *P*=0.012) and left inferior parietal (CB: 9,550±3,047 mm^3^ vs. SC: 11,769±2,064 mm^3^; *P*=0.017). Thus, the total WM volume was 13-18% lower in the CB group. Within temporal cortex, WM reductions were observed in the bilateral inferior temporal area (left: CB: 5,751±1,185 mm^3^ vs. SC: 6,927±1,190 mm^3^; *P*=0.0152; right: CB: 5,551±844 mm^3^ vs. SC: 6,704±844 mm^3^; *P*=0.0045) and right middle temporal area (CB: 5,589±842 mm^3^ vs. SC: 6,452±750 mm^3^; *P*=0.014). Finally, CB subjects had significantly reduced WM volume in right posterior cingulate cortex (CB: 4,123±1,010 mm^3^ vs. SC: 4,943±929 mm^3^; *P*=0.016), right precuneus (CB: 8,282±1,742 mm^3^ vs. SC: 9,642±1,156 mm^3^; *P*=0.013) and left medial orbitofrontal cortex (CB: 3,407±723 mm^3^ vs. SC: 4,147±464 mm^3^; *P*=0.0053).

**Figure 4:**
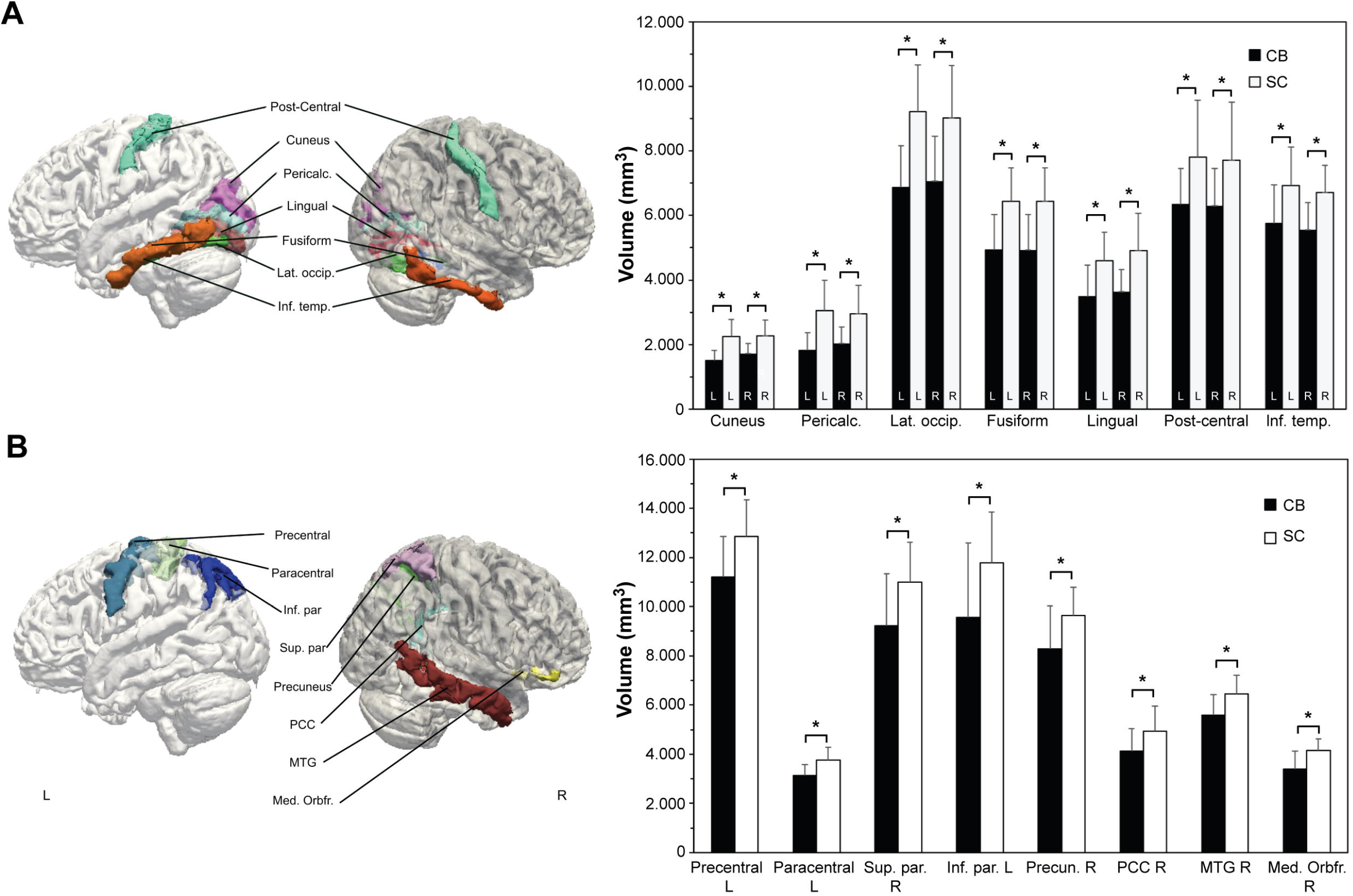
Changes in WM volume in CB subjects. ROI analysis using the Freesurfer WM atlas. The left part of the figure shows the WM ROIs with significant changes projected on a brain template. **A**. Brain white matter areas showing bilateral changes in volume. **B:** Brain white matter areas showing unilateral changes in volume. Abbreviations: Pericalc: pericalcarine; Lat. Occip: lateral occipital; Inf. Temp: inferior temporal; Sup. par: superior parietal; PCC: posterior cingulate cortex; MTG: middle temporal gyrus; Med. orbfr: medial orbitofrontal; L: left; R: right. *: P<0,05 (FDR-corrected).

### Thalamic, hippocampal and amygdalar subfields

Divisions of the thalamus showed the greatest volume differences between the CB and SC groups (Fig. 5; Table 1). The most conspicuous differences were in left (−30%) and right (- 23%) lateral geniculate nucleus. There were additional bilateral reductions in ventroposterior thalamic, centromedian, ventromedian, ventral lateral posterior, paratenial, parafascicular, central lateral, and paracentral nuclei. Other thalamic divisions were only reduced on one side, namely in the right ventral lateral anterior, ventral anterior, anteroventral, laterodorsal, reuniens and ventral anterior magnocellular nuclei, and the left mediodorsal medial magnocellular nucleus.

**Figure 5:**
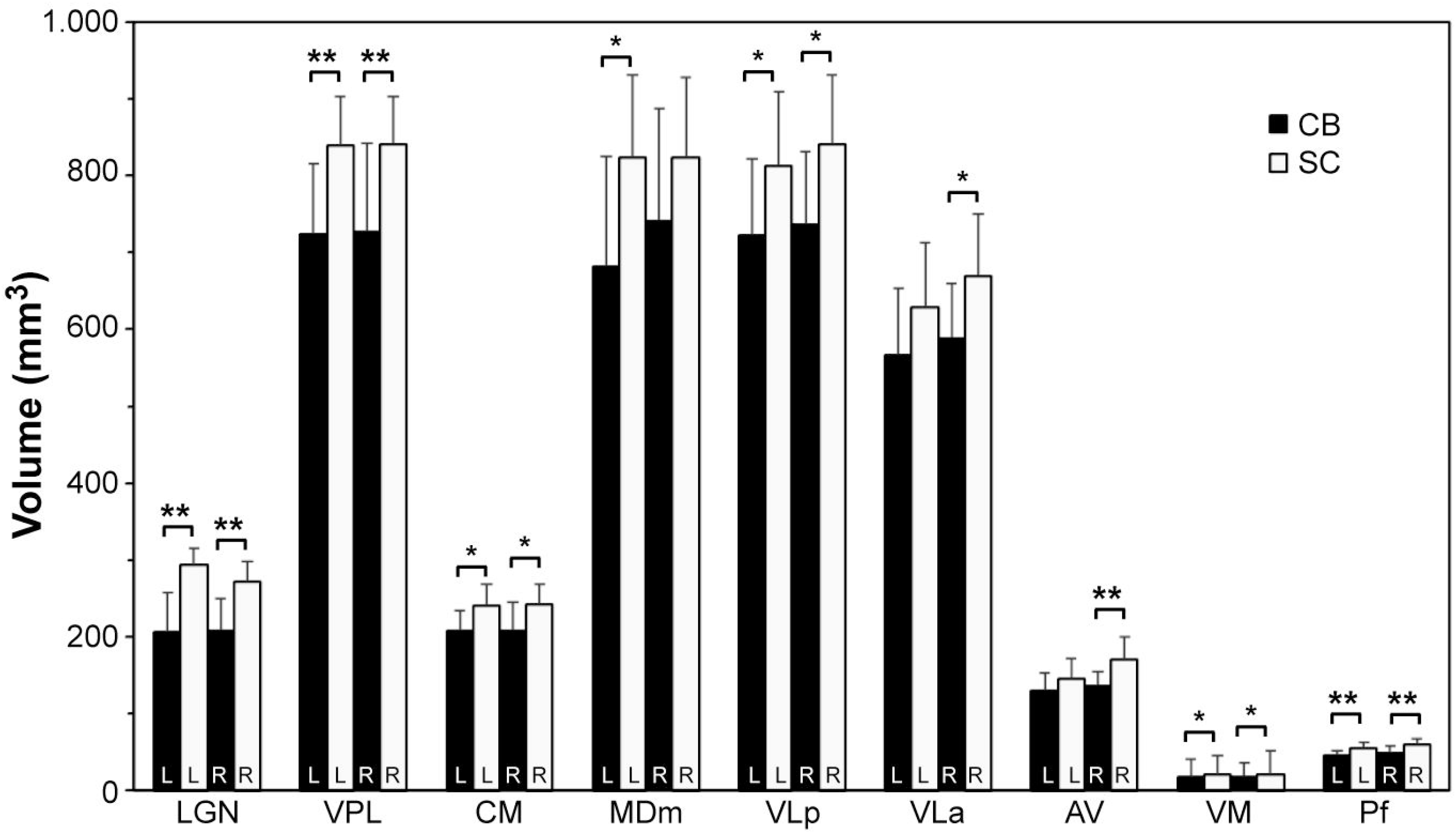
Volumetric reductions in thalamic subfields. Abbreviations: LGN: lateral geniculate nucleus; VPL: ventroposterior lateral nucleus; CM: centromedian nucleus; MDm: mediodorsal medial magnocellular nucleus; VLp: ventral lateral posterior nucleus; VLa: ventral lateral anterior nucleus; AV: anteroventral nucleus; VM: ventromedial nucleus; Pf: parafascicular nucleus; L: left; R: right. *: P<0.05 (FDR-corrected); **: P<0.01 (FDR-corrected).

**Table 1:** Volumetric measures (mean ± SD) of thalamic subfields (in mm3). All P-values are FDR-corrected.

| Group | subnucleus | Left |  |  | Right |  |  |
| --- | --- | --- | --- | --- | --- | --- | --- |
|  |  | CB | SC | P | CB | SC | P |
| Anterior |  |  |  |  |  |  |  |
|  | Anteroventral ncl. (AV) | 130 ± 23 | 145 ± 26 | NS | 137 ± 18 | 169 ± 31 | 0,001 |
| Lateral |  |  |  |  |  |  |  |
|  | Laterodorsal ncl. (LD) | 27 ± 7 | 33 ± 7 | NS | 27 ± 7 | 38 ± 38 | 0,02 |
|  | Lateral posterior ncl. (LP) | 126 ± 16 | 133 ± 17 | NS | 114 ± 17 | 133 ± 21 | 0,03 |
| Ventral |  |  |  |  |  |  |  |
|  | Ventral anterior ncl. (VA) | 405 ± 72 | 439 ± 64 | NS | 427 ± 45 | 482 ± 42 | 0,01 |
|  | Ventral anterior magnocellular ncl. (VAMc) | 30 ± 5 | 33 ± 7 | NS | 31 ± 4 | 36 ± 36 | 0,02 |
|  | Ventral lateral anterior ncl. (Vla) | 566 ± 88 | 628 ± 84 | NS | 590 ± 69 | 670 ± 81 | 0,01 |
|  | Ventral lateral posterior ncl. (VLp) | 723 ± 100 | 813 ± 97 | 0,002 | 738 ± 95 | 841 ± 90 | 0,03 |
|  | Ventroposterior lateral ncl. (VPL) | 723 ± 92 | 839 ± 65 | 0,0004 | 729 ± 113 | 840 ± 63 | 0,0009 |
|  | ventromedial ncl. (VM) | 17 ± 2 | 20 ± 2 | 0,003 | 18 ± 3 | 20 ± 3 | 0,01 |
| Intralaminar |  |  |  |  |  |  |  |
|  | Central medial ncl. (CeM) | 64 ± 13 | 76 ± 15 | NS | 66 ± 14 | 81 ± 19 | NS |
|  | Central lateral ncl. (CL) | 34 ± 5 | 43 ± 43 | 0,02 | 33 ± 5 | 46 ± 46 | 0,01 |
|  | Paracentral ncl. (Pc) | 3 ± 1 | 4 ± 4 | 0,009 | 4 ± 1 | 5 ± 5 | 0,01 |
|  | Centromedian ncl. (CM) | 208 ± 26 | 240 ± 28 | 0,004 | 209 ± 36 | 242 ± 26 | 0,005 |
|  | Parafascicular ncl. (Pf) | 46 ± 6 | 55 ± 8 | 0,01 | 50 ± 8 | 59 ± 7 | 0,003 |
| Medial |  |  |  |  |  |  |  |
|  | Paratenial ncl. (Pt) | 6 ± 1 | 7 ± 1 | 0,0006 | 6 ± 1 | 8 ± 1 | 0,001 |
|  | Medioventral ncl. (Reuniens) MV(Re) | 12 ± 3 | 15 ± 4 | NS | 12 ± 4 | 17 ± 17 | 0,02 |
|  | Mediodorsal medial magnocellular ncl. (MDm) | 682 ± 142 | 823 ± 109 | 0,008 | 724 ± 155 | 824 ± 105 | NS |
|  | Mediodorsal lateral parvocellular ncl. (MDI) | 263 ± 47 | 297 ± 42 | NS | 296 ± 48 | 306 ± 41 | NS |
| Posterior |  |  |  |  |  |  |  |
|  | Lateral geniculate ncl. (LGN) | 206 ± 50 | 294 ± 22 | 4,6E-05 | 210 ± 39 | 272 ± 27 | 0,0004 |
|  | Medial geniculate ncl. (MGN) | 96 ± 13 | 106 ± 10 | NS | 123 ± 15 | 136 ± 16 | NS |
|  | Limitans (suprageniculate) ncl. L-SG | 26 ± 6 | 29 ± 7 | NS | 26 ± 4 | 29 ± 7 | NS |
|  | Anterior pulvinar ncl. (PuA) | 224 ± 23 | 248 ± 29 | NS | 198 ± 26 | 217 ± 17 | NS |
|  | Medial pulvinar ncl. (PuM) | 1154 ± 117 | 1271 ± 153 | NS | 958 ± 111 | 1047 ± 59 | NS |
|  | Lateral pulvinar ncl. (PuL) | 231 ± 35 | 265 ± 60 | NS | 189 ± 27 | 197 ± 21 | NS |
| Total |  |  |  |  |  |  |  |
|  | Whole thalamus | 6272 ± 726 | 7174 ± 717 | 0,002 | 6128 ± 773 | 6936 ± 693 | 0,002 |

For the segmentation of the hippocampus, significant volume reductions were present only in the left presubiculum body and right CA1 area of the CB group (Table 2). When using non-FDR-corr. statistics additional volumetric reductions were found in right presubiculum body, bilateral subiculum body and head, left and right fimbria, left hippocampal tail, left granule cell layer of the dentate gyrus, and the whole right hippocampal head.

**Table 2:**
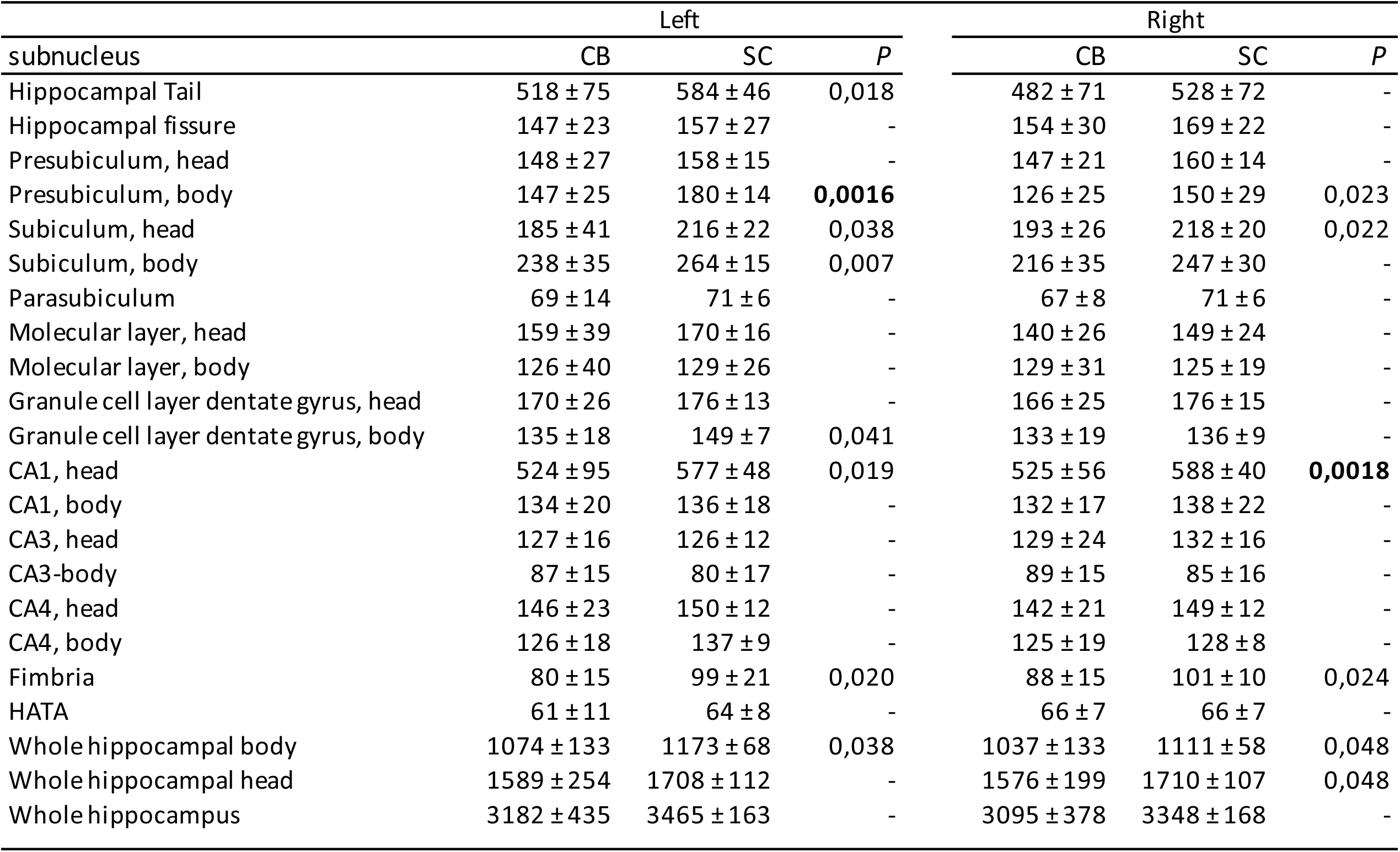
Volumetric measures (mean ± SD) of hippocampal subfields (in mm3). Values in BOLD, FDR-corrected.

| subnucleus | Left |  |  | Right |  |  |
| --- | --- | --- | --- | --- | --- | --- |
|  | CB | SC | *P* | CB | SC | *P* |
| Hippocampal Tail | 518 ± 75 | 584 ± 46 | 0,018 | 482 ± 71 | 528 ± 72 | - |
| Hippocampal fissure | 147 ± 23 | 157 ± 27 | - | 154 ± 30 | 169 ± 22 | - |
| Presubiculum, head | 148 ± 27 | 158 ± 15 | - | 147 ± 21 | 160 ± 14 | - |
| Presubiculum, body | 147 ± 25 | 180 ± 14 | **0,0016** | 126 ± 25 | 150 ± 29 | 0,023 |
| Subiculum, head | 185 ± 41 | 216 ± 22 | 0,038 | 193 ± 26 | 218 ± 20 | 0,022 |
| Subiculum, body | 238 ± 35 | 264 ± 15 | 0,007 | 216 ± 35 | 247 ± 30 | - |
| Parasubiculum | 69 ± 14 | 71 ± 6 | - | 67 ± 8 | 71 ± 6 | - |
| Molecular layer, head | 159 ± 39 | 170 ± 16 | - | 140 ± 26 | 149 ± 24 | - |
| Molecular layer, body | 126 ± 40 | 129 ± 26 | - | 129 ± 31 | 125 ± 19 | - |
| Granule cell layer dentate gyrus, head | 170 ± 26 | 176 ± 13 | - | 166 ± 25 | 176 ± 15 | - |
| Granule cell layer dentate gyrus, body | 135 ± 18 | 149 ± 7 | 0,041 | 133 ± 19 | 136 ± 9 | - |
| CA1, head | 524 ± 95 | 577 ± 48 | 0,019 | 525 ± 56 | 588 ± 40 | **0,0018** |
| CA1, body | 134 ± 20 | 136 ± 18 | - | 132 ± 17 | 138 ± 22 | - |
| CA3, head | 127 ± 16 | 126 ± 12 | - | 129 ± 24 | 132 ± 16 | - |
| CA3-body | 87 ± 15 | 80 ± 17 | - | 89 ± 15 | 85 ± 16 | - |
| CA4, head | 146 ± 23 | 150 ± 12 | - | 142 ± 21 | 149 ± 12 | - |
| CA4, body | 126 ± 18 | 137 ± 9 | - | 125 ± 19 | 128 ± 8 | - |
| Fimbria | 80 ± 15 | 99 ± 21 | 0,020 | 88 ± 15 | 101 ± 10 | 0,024 |
| HATA | 61 ± 11 | 64 ± 8 | - | 66 ± 7 | 66 ± 7 | - |
| Whole hippocampal body | 1074 ± 133 | 1173 ± 68 | 0,038 | 1037 ± 133 | 1111 ± 58 | 0,048 |
| Whole hippocampal head | 1589 ± 254 | 1708 ± 112 | - | 1576 ± 199 | 1710 ± 107 | 0,048 |
| Whole hippocampus | 3182 ± 435 | 3465 ± 163 | - | 3095 ± 378 | 3348 ± 168 | - |

There were no volumetric group differences for any of the amygdalar subfields. However, when using non-FDR thresholds, the CB group had lower volumes in in the left central nucleus and the right accessory basal nucleus (Supplementary Table 2).

### Cerebellum

The CB group had significantly lower volumes in the bilateral Crus I (right: CB: 12,870±1,520 mm^3^ vs. SC: 14,440±1,420 mm^3^; *P*=0.02; left: CB: 13,290±1,070 mm^3^ vs. SC: 14,370±920 mm^3^; P=0.02), the right lobule X (CB: 490±70 mm^3^ vs. SC: 570±70mm^3^; *P*=0.01) and lobule X bilaterally (CB: 960±160 mm^3^ vs. SC: 1,120±130 mm^3^; *P*=0.01). In addition, CB subjects had reduced cerebellar cortical thickness in right lobule V (CB: 5.09±0.18 mm vs. SC: 5.29±0.15 mm; *P*=0.008) and the bilateral lobule V (CB: 5.25±0.11 mm vs. SC: 5.37±0.13 mm; *P*=0.02). We also detected a global volumetric reduction of the right cerebellum (CB: 58,830±5,110 mm^3^ vs. SC: 65,930±6,780 mm^3^; *P*=0.007).

### Changes in cortical thickness

Overall mean neocortical thickness did not differ between the blind and sighted groups (CB: 2.28±0.70 mm vs. SC: 2.29±0.15 mm; *P*=0.62). Our ROI analysis also did not reveal significant differences in regional cortical thickness measures when using FDR-corrected statistics, but a more lenient statistical threshold (P<0.05, non-FDR-corr.) showed in the CB group lower cortical thickness in left calcarine sulcus (CB: 1.91±0.08 mm vs. SC: 1.99±0.09 mm; *P*=0.042), left postcentral sulcus (CB: 2.03±0.08 mm vs. SC: 2.14±0.09 mm; *P*=0.0027), left parieto-occipital sulcus (CB: 2.14±0.09 mm vs. SC: 2.20±0.12 mm; *P*=0.041), left anterior part of the cingulate gyrus and sulcus (CB: 2.21±0.11 mm vs. SC: 2.32±0.12 mm; *P*=0.050), and left middle anterior part of the cingulate gyrus and sulcus (CB: 2.31±0.08 mm vs. SC: 2.41 ±0.1 mm; *P*=0.019).

Next, we calculated correlations between cortical thickness measurements between regions with volumetric reductions in the group of CB participants. In the SC group, thickness in the right calcarine gyrus had strong positive correlations with thickness in the five other areas with volumetric reductions in the CB group (Fig. 6). However, in the CB group, these correlations were absent or strongly attenuated for right occipito-temporal and medial lingual gyrus (SC: R^2^=0.79 vs. CB: R^2^=0.23; *P*<0.05), right anterior collateral transverse sulcus (SC: R^2^=0.85 vs. CB: R^2^=0.004; *P*<0.05), right posterior collateral transverse sulcus (SC: R^2^=0.211 vs. CB: R^2^=0.022; *P*<0.05), right temporal pole (SC: R^2^=0.72 vs. CB: R^2^=0.04; *P*<0.05), right inferior temporal sulcus and right subcallosal gyrus (SC: R^2^=0.53 vs. CB: R^2^=0.003; *P*<0.05).

**Figure 6:**
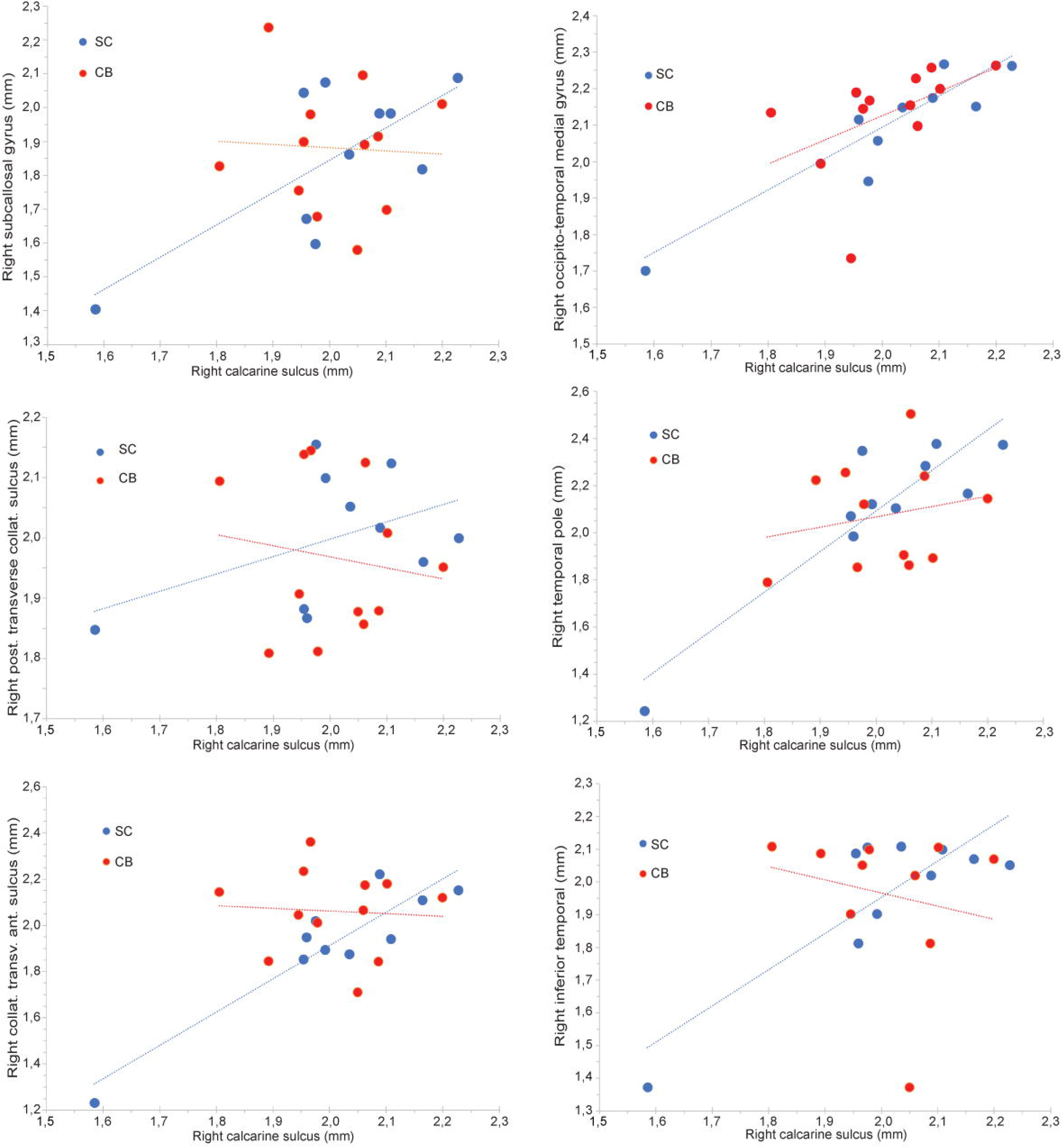
Correlations between measures of cortical thickness. Correlations between cortical thickness measures in right calcarine sulcus with thickness in give other areas with volumetric reductions (right subcallosal gyrus, occipito-temporal medial gyrus, posterior transverse collateral sulcus, temporal pole, collateral transverse anterior sulcus and inferior temporal) in the groups of congenitally blind and sighted subjects. Whereas correlations between cortical thickness measures of these areas were positive in sighted controls, the correlations were reduced or abolished in the congenitally blind participants.

### Changes in cortical curvature

For this index, group differences did not survive correction for multiple comparisons in any brain ROI region. However, using the less stringent statistical threshold of *P*<0.05, non-FDR-corr., the CB subjects showed significant alterations in cortical curvature in eight brain regions, four of which occupying occipital cortex. More specifically, the CB group had increased cortical curvature measures in left inferior occipital gyrus and sulcus (CB: 0.132±0.021 vs. SC: 0.122±0.010; *P*=0.0007, corr.), right inferior occipital gyrus and sulcus (CB: 0.136±0.024 vs. SC: 0.124±0.025; *P*=0.002), right middle occipital gyrus (CB: 0.148±0.026 vs. SC: 0.138±0.061; *P*=0.0024), and right occipital pole (CB: 0.152±0.022 vs. SC: 0.134±0.011; *P*=0.005). Outside visual areas, the CB group showed significantly decreased cortical curvature in the right intermedius primus (Jensen) sulcus (CB: 0.084±0.013 vs. SC: 0.943±0.027; *P*=0.03), but increased curvature in the orbital part of bilateral inferior frontal gyrus (left: CB: 0.158±0.025 vs. SC: 0.148±0.024; *P*=0.02; right: CB: 0.174±0.028 vs. SC: 0.160±0.024; *P*=0.04) and right anterior segment (horizontal ramus) of the lateral sulcus (CB: 0.096±0.023 vs. SC: 0.086±0.021; *P*=0.019).

## Discussion

This is the first ultra-high field MRI study on volumetric and structural brain differences in a contrast of CB individuals with sighted controls. We investigated changes in multiple parameters, including cortical volume, thickness and curvature, and WM volumes. In addition, we performed a subfield analysis of volumetric differences in thalamus, amygdala, hippocampus and cerebellum, thereby providing the most comprehensive description of structural changes in the blind brain so far. Our results show that CB is associated with important reductions in cortical and subcortical GM volumes and WM reductions, along with lesser differences in cortical thickness and curvature. The large majority of the volumetric changes were concentrated in occipital and parietal brain areas, but certain differences were also present in occipito-temporal and prefrontal cortices. We found no evidence in support of claims that congenital blindness may lead to adaptive or compensatory increases in the volumes of non-visual brain areas, as reported by some authors.^15,16,22-25^

### Changes in global measures

Compared to sighted controls, CB subjects had lower mean volumes for total cerebral cortex, GM, WM, and brainstem, with reductions in the range of 12 - 15%. This is in line with some earlier reports of a global reduction in overall measures of brain GM and WM volumes of CB individuals.^6,9,11,26,27^ We also report volume reductions in subcortical structures, notably in the nucleus accumbens, which has not hitherto been noted in CB individuals, although a recent 7T MRI study reported a volume reduction of the right accumbens in individuals with late-onset blindness (Jonak et al., 2020).^28^ The nucleus accumbens receives afferent input from mesolimbic dopaminergic projections, as well as direct input from prefrontal, somato-motor, anterior cingulate and subgenual cingulate cortices, entorhinal and perirhinal cortex, hippocampus, thalamus and amygdala. The accumbens sends projections back to most of these areas and also to the basal ganglia.^29,30^ Along with the rest of the ventral striatum, the nucleus accumbens is involved in motivational and emotional processes and is implicated in a multitude of (neuro)psychiatric and neurological disorders.^31^ The nucleus accumbens also plays a role in the perception of pain and pain regulation.^32,33^ We have shown that CB subjects are more sensitive to pain^34,35^ and are more vulnerable to the psychological context in which a painful stimulus is presented.^36^ The present findings of significant volumetric reduction of the nucleus accumbens in CB subjects may hence relate to their known alterations in pain perception and regulation.

Our data also revealed volume reductions of the caudate nucleus and globus pallidus, the first such report for globus pallidus in CB subjects. One earlier study reported a GM increase of the right globus pallidus in CB subjects born at term, but not in CB subjects born at pre-term.^37^ The globus pallidus receives major input from the striatum, the subthalamic nucleus, and several cerebral cortical regions, and project to the thalamus, which relays further to prefrontal and other cortical regions.^38^ Although its principal role is to control conscious and proprioceptive movements, the globus pallidus has other functions involved in cognition and motivation. Considering the role of vision in motor planning, we propose that volumetric reduction of the globus pallidus relates to the difficulties for the blind in fine motor, balance and trunk control.^39^

In line with our earlier findings^6^, the caudate nucleus volume was also reduced in CB subjects. The caudate is part of the dorsal striatum, which, together with the hippocampus, is integral to memory as pertains to spatial navigation. Whereas the hippocampus is involved in allocentric forms of navigation, the caudate nucleus is implicated in egocentric navigation, and in building stimulus-response strategies.^40^ Route-learning, which involves a sequence of stimulus-response associations, activates the caudate nucleus^41^ and its proficiency correlates with caudate volume.^42^ The caudate nucleus also engages in movement planning and execution, memory, learning, reward, motivation, and emotion.^43^ Efferent projections from the caudate nucleus innervate the hippocampus, globus pallidus and thalamus, and its afferent arise in broad inputs from visual area TE in temporal cortex.^44^ Single unit recordings in the macaque caudate showed cellular responses to visual stimuli^45^ and to eye movements, and revealed an association between neuronal firing rate to the reward value of the stimulus.^46^ The caudate nucleus is also a key node in the closed-loop visual cortico-corticostriatal pathway.^47^ Its volumetric reduction in CB individuals may thus related to their problems in route learning^48^ and certain higher cognitive functions.^43^

The brainstem and ventral diencephalon were also smaller in our group of CB subjects. In the Freesurfer segmentation, “ventral diencephalon” refers to a conglomerate of different nuclei and surrounding WM, including the hypothalamus, the mammillary bodies, the subthalamic nucleus, lateral and medial geniculate nucleus, the red nucleus and substantia nigra. The volumetric reduction of this zone in our CB group could hence arise from reduced volume in any or all of its constituent nuclei. The suprachiasmatic nucleus, located in the anterior hypothalamus, receives direct visual input via the retino-hypothalamic tract^49^ and is involved in the regulation of circadian rhythms. That CB subjects have alterations in their circadian rhythms^50,51^ may conceivably relate to the absent visual input to the suprachiasmatic nucleus.

### Alterations in cortical grey matter volume

The CB group had reduced cortical volume in the occipital, occipito-temporal, parietal, and prefrontal lobes. Occipital (visual) areas with reduced volume included the cuneus and the occipital pole, and the posterior transverse collateral sulcus and the adjacent medial occipito-temporal and lingual sulci in the inferior temporal cortex. Here, the volumetric reductions were among the most pronounced, exceeding 30% for the cuneus and posterior transverse collateral sulcus relative to the control group. The volumetric reductions were smallest in the more anterior situated medial occipito-temporal and lingual sulci, suggesting a gradient in which primary visual cortex is more strongly affected than higher order visual areas. These results are in line with earlier reports of volumetric reductions in the cuneus, calcarine and pericalcarine sulcus, and the lingual and fusiform gyri.^6,11,25,37,52-56^

Interestingly, we found no sign of volumetric reductions in more dorsally situated visual areas, which may indicate that GM areas within the ventral visual stream are more prone than the dorsal stream to effects of lifelong visual deprivation. This conjecture is in line with our previous findings that blindness selectively alters the microstructure of the ventral but not the dorsal visual stream.^57^ That finding might reflect differences in the developmental time course of the maturation of the ventral and dorsal visual streams in the human brain. Although functional maturity of the dorsal stream precedes that of the ventral stream^58^, WM structural maturation is slower in the ventral stream.^59^

The left central and precentral sulci were also smaller in CB subjects. Similar trends were seen for the homologous areas in the right hemisphere but did not survive correction for multiple comparisons. This first report of a volumetric reduction in the somatosensory cortex of CB individuals is in agreement with the report of a volumetric reduction of the primary somatosensory cortex in congenitally anophthalmic mice.^60^ Our data are also in line with our earlier magnetoencephalography data showing that CB subjects have reduced directed functional connectivity from primary somatosensory cortex to thalamus in the beta frequency range.^61^ Task-related beta-oscillations are likely to convey top–down signaling across brain networks and systems.^62^ Our morphometric data therefore may constitute a neuroanatomical correlate for the hypothesized reduction of feedback processes in CB individuals in the context of somatosensory stimulation.^61^ The volumetric reduction of the somatosensory cortex may also be related to the smaller volume of the ventroposterior thalamic nucleus in the CB subjects.

Finally, CB participants had reduced volume of the left superior frontal and the right subcallosal gyri. The superior frontal sulcus contains the frontal eye fields, which are involved in the control smooth pursuit and saccadic eye movement.^63^ Two previous studies have reported anatomic changes in the frontal eye fields area in blind subjects. Pan and colleagues^55^ showed a cortical thinning in this area, whereas Voss and Zatorre^12^ showed a reduced functional connectivity between the frontal eye fields and the calcarine sulcus. On the other hand, the subcallosal gyrus is part of a network linking the orbitofrontal and medial prefrontal cortices and the agranular insula cortex with the limbic system, thalamus, hypothalamus and brainstem. Metabolic rate is reduced in the subcallosal gyrus in patients suffering from major depression, which can be rectified by antidepressant medication or deep brain stimulation of this area.^64^ The volumetric reduction in the subcallosal gyrus is in line with reports of increased depressive symptoms in those with visual impairment^65^ and with the importance of light perception on mood in general.^66^

### Alterations in white matter

The WM reductions in the CB group largely match those observed in cortical GM, being concentrated mainly in occipital, occipito-temporal and parietal areas. More specifically, reduced WM volume was detected in the cuneus, occipital pole, the adjoining posterior transverse collateral sulcus, as well as in the more anterior medial occipito-temporal and lingual gyri. Much as for cortical GM, these WM reductions were substantial, ranging from 30 to 40% in cuneus and peri-calcarine area 20 to 25% in other visual areas, thus matching the GM reduction in the lateral geniculate nucleus. This implies a causal cascade of transsynaptic loss within the retina-geniculate-striatal pathway. A few earlier studies also reported WM reductions in the cuneus ^6,26,48^ and lingual gyrus.^26,48,67^ Our finding of WM decrease in somatosensory cortex is in line with the concomitant GM reduction in this area, but is at odds with results of an earlier study reporting significant WM *increases* in this area in CB individuals.^25^ Our CB subjects also showed volumetric WM loss in the superior frontal and the subcallosal gyri, two areas that also had GM loss.

The present data also confirm our previous reports of WM volumetric alterations in the optic chiasm and the splenium of the corpus callosum.^68-70^ Callosal fibers of the splenium convey visual information between the hemispheres, thereby connecting large areas of the left and right occipito-temporal and occipito-parietal cortices. Of note, the 18% lower WM volume in the splenium matches that in our earlier reports.^68,69^ The more than 80% of the splenium remains in the CB raises the question of what type of information these fibers convey. A vast number of studies have shown that congenital blindness in humans leads to a reorganization of the brain through a process of cross-modal plasticity.^5^ In addition, congenital blindness is associated with important alterations in the functional connectivity of the occipital cortex.^61,70- 73^ Together, these findings may explain the resilience of volumetric loss in the splenium volume and visual cortical areas of the CB.

### Alterations in cerebellum

The cerebellum showed several volumetric alterations in CB subjects, in keeping with an earlier report.^74^ However, that study neither distinguished between GM and WM changes, nor did it investigate specific lobular changes. We now report that the relative WM reduction exceeds by nearly two-fold that for GM in cerebellum of the CB. This is in contrast to our neocortical findings, showing relatively greater reductions in GM than in WM. In sighted individuals, the integrity of cerebellar WM plays an important role in reading skills.^75,76^ The pronounced loss of cerebellar WM in blind individuals may hence reflect their use of somatosensory cues for reading. Alternately, the present findings may be a consequence of pre-term birth^77,78^ which had caused retinopathy of more than half of our blinds group.

Besides the overall reduction in cerebellar GM and WM volumes, our data revealed a localized volumetric reduction in Crus I, which is in the supramodal zone of the cerebellum, having strong reciprocal projections with prefrontal cortex (BA 46) and posterior parietal cortex.^79^ As such, Crus I plays a strong role in spatial navigation. Activation studies show that, whereas the left Crus I, in combination with the right hippocampus and the medial parietal cortex, is involved in place-based navigation, the right lobule Crus I, in conjunction with left hippocampus and the medial prefrontal cortex, is implicated in sequential egocentric representation.^80^ Our finding of a volumetric reduction in Crus I is therefore in line with the well-known association of impairments in spatial navigation capabilities with congenital blindess.^81^ However, other plausible interpretations include the involvement of Crus I in language and executive functions such as working memory and planning, ^82^ which are affected in CB individuals.^5^ In addition, a recent 7T fMRI study showed that the cerebellum contains whole-body somatotopic maps in which the eyes are represented in lobules VI, VIIb and Crus I.^83^ Finally, Lobule X was also reduced in volume in the present group; this relatively small lobule is involved in emotion processing and in vestibular control.^84^

### Group differences in cortical thickness and gyrification

In contrast to the robust changes in cortical volume, there were relatively few group differences in cortical thickness. Only when using uncorrected P-values, we identified some cortical areas with reduced cortical thickness in the congenitally blind group. These included two areas in the occipital cortex, the left calcarine and parieto-occipital sulci, the bilateral post-central sulcus, and a region in the left mid-anterior cingulate cortex. There was no trend towards increased cortical thickness in the peri-calcarine area.

The literature on changes in cortical thickness in congenital blindness is not very consistent. Of 14 studies, eight reported an increase in cortical thickness in the peri-calcarine area in CB individuals.^10,12,52,54,56,85,86,87^ The remaining studies reported no changes in this area, as in our present investigation. Two studies reported a decrease in cortical thickness in the post-central sulcus in CB subjects^9,13^, whereas three others reported increased thickness of this area^10,14,87^. Finally, three papers mentioned increased cortical thickness in the anterior cingulate cortex in CB individuals^9,10,14^, whereas the other studies did not mention changes in this area. The reason for this discrepancy is unclear; our present use of a 7T magnet with acquisitions at submillimeter resolution should in principle have given more reliable estimates of cortical thickness compared to older studies

Although we found scant evidence for robust changes in cortical thickness in CB subjects, we did observe certain alterations in covariance measures of cortical thickness. Whereas cortical thickness of the right calcarine sulcus correlated positively with thickness of six other ROIs showing volumetric changes in the group of blind subjects, this correlation was strongly reduced or absent in the CB group, suggesting alterations in the connectivity of the calcarine sulcus with these areass.

Our structural analysis revealed four regions within the occipital cortex showing non-significant trends (significant at non-FDR-corrected level) for increased gyrification in CB. These regions included the inferior occipital gyrus and sulcus (bilateral), the right occipital pole and middle occipital gyrus, regions which also showed decreased volume in the CB group. Cortical folding represents an indirect measure of brain structural connectivity that depends on the thickness ratio between outer and inner cortical layers.^88,89^ Consequently, a higher folding complexity (i.e., more curvature) is indicative of increased cortico-cortical connectivity and relatively reduced descending projections.^90,91^ Studies conducted in a rhesus macaque model of congenital blindness showed that retrograde tracers injected in the visually deprived cortex reveal a relatively increased local, intrinsic connectivity characteristic of higher order areas.^92^ We therefore hypothesized that the absence of visual input from birth promotes local cortico-cortical connectivity^93^, manifesting in greater gyrification of the cortical surface within this region. Notably, such changes could occur in conjunction with a reduction in cortical volume.

### Alterations in thalamic subfields

The thalamus showed significant volumetric reductions in several subfields, most prominently in the left (−30%) and right (−23%) lateral geniculate nuclei. This may be unsurprising, given their role as principal recipients of innervation by the optic nerve and tract. This present result is in line with previous findings from our group, showing a similar volumetric reduction in the lateral geniculate nuclei in CB subjects, in which we used an in-house segmentation tool to subdivide the thalamus.^69,93^ Also in line with those earlier reports, we found no evidence for volumetric alterations of the adjacent medial geniculate nucleus, the principal thalamic relay for auditory input. The VPL, which is the main relay nucleus for tactile and thermal sensory inputs, also showed a robust volumetric bilateral volume decrease in the CB group. That result fits with the volumetric reduction in the (sensory) post-central gyrus of the cerebral cortex. However, studies in mice enucleated at birth showed preservation of the volume of the VPL^60^ and of its projections to SI.^94^ Additional reductions were found in the ventromedial, ventrolateral, mediodorsal and centromedial thalamic nuclei. The ventromedial and ventrolateral nuclei comprise the ventral motor nuclei of the thalamus, which conveys motor-related information to the prefrontal cortex. However, recent studies have shown that these nuclei also have reciprocal connections with the prefrontal cortex and contribute decision making, an executive function^95^ and in the regulation of cortical arousal.^96^ The centromedian nucleus projects to the putamen and receives inputs from the globus pallidus, the superior colliculus and the reticular formation; it is involved in cognition and sensory-motor integration.^97^ Finally, the mediodorsal nucleus has reciprocal connections with the dorsolateral, ventromedial and orbitofrontal cortex, and the anterior cingulate cortex, via which it plays an important role in cognition.^98^

Overall, these volumetric changes in the thalamus are in line with diffusion imaging data from our lab showing microstructural changes in the temporal, sensory, and frontal subdivisions of the thalamus in CB subjects.^99^ However, present results differ from another MRI study in which the volumes of thalamic relay nuclei directly involved in motor and somatosensory processing were unaffected by visual deprivation.^93^ Some of these differences may be attributable to our use of a 7T magnet and a different segmentation tool.

Although the hippocampus did not show an overall volume reduction in volume in the CB participants, there were focal reductions of the presubiculum body and CA1 The presubiculum receives integrated information from central and peripheral visual inputs via the parieto-medial temporal pathway.^100^ Originating in the caudal inferior parietal lobule, this pathway and conveys visuospatial information from parietal regions to specific regions in the medial temporal lobe.^101^ The presubiculum famously contains grid, border and head direction cells^102,103^ and plays a role in scene-based cognition.^101^ We also saw volumetric reduction in the right CA1, which plays an important role in the formation, consolidation, and retrieval of hippocampal-dependent memories. In healthy individuals, CA1 volume correlates positively with episodic memory retrieval ^104^ and focal lesions to the CA1 region disturb autobiographical memory.^105^ When using non-FDR-corrected statistics, we also found a volumetric reduction in the hippocampal tail, in line with our earlier findings.^7^ The posterior (dorsal) hippocampus plays an important role in spatial navigation in normal sighted individuals.^106,107^ The subiculum also showed a bilateral volumetric reduction when using non-FDR-corrected statistics. Situated between CA1 and the presubiculum, the subiculum is closely linked to the hippocampus proper via dense inputs from CA1, it is functionally independent from the rest of the hippocampus.^108^ The subiculum sends projections to a large number of cortical and subcortical brain areas via the fornix; it is involved in episodic memory, and especially spatial working memory.^108^

We saw no significant volumetric group differences for any of the amygdalar subfields. This was quite surprising, since the amygdala receives multiple visual inputs from subcortical pathways via the superior colliculus and the thalamic pulvinar, and also via a pulvinar-superior temporal sulcus connection. The amygdala also receives visual inputs from the retino-geniculate pathway and through the ventral stream, though potentially also through a dorsal V1-superior temporal sulcus - amygdala pathway. Nonetheless, the amygdala volumes are seemingly unaffected by CB.

### Study limitations

The current study included a total of 12 CB subjects which is a relatively small sample size for brain morphometric analysis. The small sample size reflects the difficulty of finding CB subjects in modern Western societies. Another limiting factor is the lack of behavioral correlates in this study. Next, our results are derived from atlas-based segmentations which are often less precise than manually drawn regions-of-interest. Finally, we did not measure Braille reading performance, tactile or auditory thresholds, spatial navigation, pain perception, or any psychological trait in our sample of subjects. Future studies should consider collecting more behavioral and neuropsychological data in order to explore purported correlations with the brain morphometric changes.

## Conclusion

We used ultra-high field MRI at submillimeter resolution to provide the most detailed description of brain structural brain changes in congenital blindness. Our results show widespread morphometric changes of the GM and underlying WM of occipital and parietal brain areas, with additional changes in occipito-temporal and prefrontal cortices. Cortical thickness was seemingly preserved. Our data also revealed hitherto unreported volumetric reductions in several subcortical areas and in divisions of the cerebellum. We found no evidence in support of the theory that congenital blindness causes adaptive structural alterations leading to increased volume in non-visual brain areas. This mismatch between seeming morphological atrophy and superior performance in various non-visual tasks, as reported in many behavioral studies, is a conundrum waiting to be solved.

## Abbreviations

CB: congenital blindness
GM: grey matter
SC: sighted controls
WM: white matter

## Acknowledgments

The authors would like to thank professor Paul Cumming (University of Bern, Austria) for critical reading of the manuscript.

## Supplementary material

Supplementary material is available at *Brain* online

