## Supplemental Table 1 for "Brain morphometric changes in congenitally blind subjects: a 7 Tesla MRI study"

**Supplementary Table 1: demographics blind participants**

| Sex | Age | Cause | Onset | Residual vision |
|---|---|---|---|---|
| M | 45 | Retinopathy of prematurity | at birth | none |
| M | 45 | Leber's congenital amaurosis | at birth | none |
| M | 45 | Retinopathy of prematurity | at birth | none |
| M | 45 | Retinopathy of prematurity | at birth | none |
| F | 39 | Leber's congenital amaurosis | at birth | minimal light sensitivity |
| F | 34 | unknown | at birth | none |
| F | 53 | Retinopathy of prematurity | at birth | none |
| M | 47 | Retinoblastoma | at birth | none |
| F | 62 | Retinopathy of prematurity | at birth | none |
| M | 53 | Retinopathy of prematurity | at birth | minimal light sensitivity |
| M | 38 | Leber's congenital amaurosis | at birth | minimal light sensitivity |
| F | 29 | Retinopathy of prematurity | 2 yrs | none |
