## Supplemental Table 2 for "Brain morphometric changes in congenitally blind subjects: a 7 Tesla MRI study"

**Supplementary Table 2: Volumetric measures amygdalar subfields (in mm3).** Table shows non-FDR-corrected P-values

| | Left | | | Right | | |
|---|---|---|---|---|---|---|
| subnucleus | CB | SC | *P* | CB | SC | *P* |
| Lateral nucleus | 620 ± 88 | 666 ± 47 | - | 612 ± 81 | 670 ± 78 | - |
| Basal nucleus | 425 ± 71 | 453 ± 32 | - | 423 ± 50 | 461 ± 51 | - |
| Accessory basal nucleus | 253 ± 49 | 270 ± 20 | - | 260 ± 38 | 284 ± 24 | 0,045 |
| Anterior amygdaloid area (AAA) | 51 ± 9 | 54 ± 5 | - | 57 ± 4 | 59 ± 5 | - |
| Central nucleus | 43 ± 11 | 46 ± 5 | - | 45 ± 7 | 52 ± 3 | 0,010 |
| Medial nucleus | 19 ± 5 | 21 ± 3 | - | 23 ± 5 | 25 ± 4 | - |
| Cortical nucleus | 25 ± 6 | 24 ± 2 | - | 26 ± 4 | 27 ± 3 | - |
| Corticoamygdaloid transition zone | 189 ± 33 | 192 ± 21 | - | 197 ± 29 | 201 ± 17 | - |
| Paralaminar nucleus | 49 ± 8 | 50 ± 5 | - | 48 ± 5 | 51 ± 5 | - |
| Whole amygdala | 1674 ± 266 | 1776 ± 102 | - | 1692 ± 209 | 1829 ± 172 | - |
